# Trait Absorption Amplifies the Path to Spatial Presence in Highly Immersive Virtual Reality: Attentional Mediation and Dose-Response Effects

**DOI:** 10.64898/2026.03.03.709394

**Authors:** Hannah R. Hayes, Carlo Campagnoli

## Abstract

Virtual Reality (VR) applications depend on eliciting spatial presence, the subjective experience of being physically located within a virtual environment. Although individual differences have long been theorised to contribute to this experience, their role in highly immersive VR systems remains contested. The present study investigated whether trait absorption predicts spatial presence and whether this relationship is mediated by attention allocation. Seventy participants (44 female, 26 male; M age = 22.90, SD = 4.88) completed a 6-minute VR session using a Meta Quest 3 Head-Mounted Display and validated self-report measures of trait absorption (Tellegen Absorption Scale), attention allocation, and spatial presence (MEC-Spatial Presence Questionnaire). Path analysis confirmed a significant, complete mediation pathway: trait absorption positively predicted attention allocation (β = 0.27, p = .013), which in turn strongly predicted spatial presence (β = 0.54, p < .001). The direct path from absorption to spatial presence was non-significant (β = 0.11, p = .325), indicating complete mediation. The indirect effect was significant (β = 0.15; 95% BCa CI [0.025, 0.291]). The model explained a sizeable 33.8% of the variance in spatial presence (Cohen’s f² = 0.51). Post-hoc dose-response analysis revealed that trait absorption acts as a cognitive amplifier: the strength of the attention–presence relationship tripled from low-absorption (β = 0.33, R² = .15) to high-absorption individuals (β = 1.00, R² = .56). These findings demonstrate that individual differences remain important in highly immersive VR by modulating the effectiveness of attentional focus, offering promising directions for tailoring VR interventions.

## 1. Introduction

### 1.1 The Centrality of Presence in Virtual Reality

Virtual Reality (VR) technology has advanced considerably in the last decade, becoming more accessible and sophisticated than ever before (Riboni et al., 2020). Its capacity to generate immersive, three-dimensional virtual environments (VEs) provides powerful solutions to practical barriers in clinical therapy and education (Emmelkamp & Meyerbröker, 2021; Cipresso et al., 2018). Technologies such as Head-Mounted Displays (HMDs) immerse the user’s senses in a VE, encouraging exclusion from the physical world (Abbas et al., 2023; Fu et al., 2023) and often eliciting thoughts and behaviours as if the VE were actual reality, even when the user is fully aware it is not (Sanchez-Vives & Slater, 2005). This psychological state, widely referred to as ‘presence’ (the feeling or illusion of ‘being there’), is considered the ultimate goal of VR development (Weech et al., 2019; Lombard & Ditton, 1997; Slater, 2003; Schubert, 2009).

The success of many VR applications, from exposure therapy for anxiety disorders to surgical simulation, hinges on the successful elicitation of presence (Carl et al., 2019; De Leo et al., 2014; Riva et al., 2019; Valmaggia et al., 2016). Despite its importance, the field has struggled with a lack of conceptual clarity and theoretical consensus, a well-documented problem that has produced a proliferation of measurement tools often conflating the experience of presence with its antecedents (Lombard & Ditton, 1997; Hein et al., 2018; Souza et al., 2021; Lombard & Jones, 2015). Many popular questionnaires include items related to technology quality or user traits, which, while potentially influential, are argued to reflect correlates of presence rather than the core construct itself (Slater, 1999; Böcking et al., 2004; Nannipieri, 2022). Many measures also tie presence to media-related factors, fundamentally undermining its status as a psychological phenomenon that can also occur outside of technology, such as during reading (Schubert & Crusius, 2002). The variability across measures hinders cross-study comparison and the application of larger-scale analytical methods needed to address the lack of conceptual clarity (Grassini & Laumann, 2020; Kukshinov et al., 2025; Cummings & Bailenson, 2016).

### 1.2 Spatial Presence: A Theoretically Grounded Approach

Given these challenges, spatial presence offers a more parsimonious and theoretically grounded alternative (Laarni et al., 2015; Wirth et al., 2007). This concept is founded in the theory that the individual’s experience of presence in VR reflects the same internal processes that generate spatial awareness in the real world (Biocca, 1997; Loomis, 1992). Spatial presence comprises two dimensions essential for constructing a mental model of one’s spatial environment: perceived self-location within the environment and an awareness of the possible actions available within it (Schubert, 2009; Glenburg, 1997; Zahorik & Jenison, 1998). These two conditional requirements for perceiving the world have gained wide agreement as the direct components of spatial presence in both physical and mediated environments (Hartmann et al., 2015; Schubert et al., 2001; Wirth et al., 2007). Measuring spatial presence through these components enables the separation of the outcome from debated antecedents, facilitating clearer interpretation of investigated influences.

For spatial presence to emerge, the sensory input provided by the VE must be rich and perceptually coupled to the user’s expectations and sensorimotor models (Lopez et al., 2015; Slater et al., 1995; Regenbrecht & Schubert, 2002). Bottom-up, automatic processing incited by sensory features of the VE interacts with the user’s traits and expectations, alongside higher-order top-down processing (Hartmann et al., 2015; Hofer et al., 2012; Schubert, 2009; Wirth et al., 2007). The sensory affordances of the VR system, specifically its level of immersion, facilitate bottom-up processing by capturing attention within the VE and encouraging exclusion from the real world (Slater & Wilbur, 1997). Meta-analytical findings indicate a medium-sized effect of immersion on spatial presence (Cummings & Bailenson, 2016). However, some authors have argued that the influence of individual user factors is diminished in highly immersive VEs (Sas & O’Hare, 2003; Uz-Bilgin & Thompson, 2022), and given the rapid advancement of VR technology (Li & Lee, 2022), it remains unclear how top-down, user-driven processes contribute to spatial presence in the context of modern, highly immersive VR.

### 1.3 Individual Differences: Trait Absorption and Attention

One of the most theoretically promising individual difference variables for explaining presence is trait absorption, defined as “a disposition for having episodes of ‘total’ attention that fully engage one’s representational (i.e., perceptual, enactive, imaginative, and ideational) resources” (Tellegen & Atkinson, 1974, p. 268). It is proposed to support top-down attentional processing that reinforces spatial presence, either by increasing a user’s propensity for focused attention toward the VE or by enhancing motivation to become involved in it (Wirth et al., 2007). Both pathways implicate attentional resources in the formation of presence. Spatial presence models consistently emphasise that attention must be directed away from the real world and toward the VE to form a coherent mental model of the mediated space (Hofer et al., 2012; Schubert, 2009; Hartmann et al., 2015). Accordingly, it has been posited that trait absorption may facilitate spatial presence through this attentional mechanism, specifically by increasing the user’s capacity or tendency to allocate attention selectively to the VE (Kober & Neuper, 2012; Sacau et al., 2005).

However, empirical support for this pathway has been mixed. In lower-immersive systems, absorption is identified as a predictor of presence (Kober & Neuper, 2012; Sacau et al., 2005; Sas & O’Hare, 2003), albeit with some discrepant findings (Chiquet et al., 2023). Null findings in higher-immersive VEs (Murray et al., 2007; Schuemie et al., 2005; Chiquet et al., 2023) seemingly support claims that the attentional mechanism of absorption becomes negligible in the presence of bottom-up processing driven by highly immersive technology (Sas & O’Hare, 2003). However, such conclusions are complicated by the use of differing measures of both presence and absorption across studies, making it insufficient to attribute discrepant findings solely to the immersive features of the VE (Sacau et al., 2008; Hein et al., 2018). For example, while Wirth et al. (2012) found absorption to be highly predictive of spatial presence using the Tellegen Absorption Scale (TAS) and the MEC-Spatial Presence Questionnaire in a lower-immersive context, Chiquet et al. (2023) reported null findings in a highly immersive VE using a pictorial scale for spatial presence and the absorption subscale of the Immersive Tendencies Questionnaire. Concerns around this comparison are amplified by evidence questioning the psychometric adequacy of the ITQ absorption subscale (Parsons et al., 2015). These differences raise the possibility that observed discrepancies are attributable to inconsistencies in measurement, rather than immersion level.

Although attention is widely theorised as the mechanism through which absorption operates on spatial presence (Wirth et al., 2007; Hartmann et al., 2015), this attentional mechanism has remained largely untested, particularly in highly immersive VEs. The present investigation seeks to directly test this pathway by measuring attention allocation separately from, and alongside, trait absorption.

### 1.4 The Present Study

The present study aimed to test the relationships between trait absorption, attention allocation, and spatial presence in a highly immersive, HMD-generated VE. Using validated measures and a sample powered to detect medium-sized effects, this study addresses key methodological limitations of prior work. We hypothesised that (H1) trait absorption will positively predict spatial presence, and that (H2) this relationship will be mediated by attention allocation. Depressive symptomatology was also measured to explore its potential moderating role on the attention–presence pathway. By isolating spatial presence from both technology- and user-related antecedents, the present investigation seeks to clarify the mechanism through which trait absorption influences presence in virtual reality.

## 2. Method

### 2.1 Participants and Design

An opportunity sample of university undergraduates was recruited via social media. After data cleaning, the final sample consisted of 70 participants (44 female, 26 male; M age = 22.90, SD = 4.88, range = 19–49). All participants reported normal or corrected-to-normal vision. Exclusion criteria included a history of cybersickness, given its negative impact on presence and participant comfort (Weech et al., 2019). Participants’ previous VR experience ranged from no previous use (n = 34), rare use (n = 33), to monthly (n = 2) or weekly use (n = 1). Informed consent was obtained from all participants prior to participation. The study employed a correlational, cross-sectional design. Ethical approval was granted by the University of Leeds School of Psychology Research Ethics Committee (PSCETHS-1384).

An a priori power analysis indicated that, to detect a medium-sized effect (f² = 0.15) in a multiple regression model with two predictors at α = .05, a sample of approximately 57 participants would be required to achieve 80% power. The final sample of 70 participants was therefore adequately powered for the primary analyses.

### 2.2 Measures

#### Trait absorption

The Tellegen Absorption Scale (TAS; Tellegen & Atkinson, 1974) was used to measure trait absorption. The scale consists of 34 True/False items summed to produce a total score (possible range: 0–34), with higher scores indicating greater trait absorption. The TAS has demonstrated excellent reliability in immersive research (Parsons et al., 2015), and the Kuder–Richardson 20 coefficient in the present sample was .90.

#### Spatial presence

The MEC-Spatial Presence Questionnaire (MEC-SPQ; Vorderer et al., 2004) was used to measure spatial presence, employing the 6-item self-location and 6-item possible-actions subscales. Items are scored on a 5-point Likert scale ranging from 1 (‘I do not agree at all’) to 5 (‘I fully agree’). Scores were summed across both subscales to produce a composite spatial presence score (possible range: 12–60), with higher scores indicating greater spatial presence. Internal consistency of the self-location (α = .93), possible-actions (α = .90), and combined spatial presence scale (α = .93) was excellent.

#### Attention allocation

The 6-item attention allocation subscale of the MEC-SPQ (Vorderer et al., 2004) was used to measure the degree to which participants directed their attention toward the VE. Items use the same 5-point Likert scale (possible range: 6–30), with higher scores indicating greater attention allocation. Internal consistency was excellent (α = .91). One participant had a single missing item on this scale, which was imputed with the participant’s item-level median prior to computing the total score.

#### Depressive symptomatology (exploratory)

To explore whether depressive symptomatology moderated the attention–presence pathway, the Beck Depression Inventory–II (BDI-II; Beck et al., 1996) was administered, consisting of 20 items (one item was removed for ethical reasons) on a 0–3 scale. In addition, the Brief Ruminative Response Scale (BRRS; Topper et al., 2014), a 5-item measure scored on a 4-point Likert scale, was administered to capture ruminative cognitive style. The BRRS was rescaled to a 0–3 range to align with the BDI-II, and scores from both instruments were summed to create a composite depression variable.

### 2.3 Apparatus and Stimuli

A Meta Quest 3 HMD was used, featuring a per-eye resolution of 2064 × 2208 pixels and a 90 Hz refresh rate (Meta, 2023), along with two Touch Plus controllers. The VE was a realistic and emotionally neutral simulation of Leonardo da Vinci’s Workshop (Stega, 2023), featuring several interactive activities. This environment was selected to minimise confounding effects of emotional salience on spatial presence (Wirth et al., 2012) and to provide a plausible, realistic context supportive of spatial presence formation (Wirth et al., 2007). Participants were confined to a painting activity within the workshop, where they could use paintbrushes and pigments to paint on a canvas. Due to space constraints, participants’ physical movement in the VE was produced by their own body movements rather than controller-based locomotion, as physical movement is known to enhance spatial presence (Slater et al., 1995). The participant’s viewing scene was streamed to a laptop via USB-C cable for researcher monitoring.

### 2.4 Procedure

Participants completed the pre-exposure questionnaire online via Qualtrics (Qualtrics, 2025), including informed consent and measures for trait absorption and depression. This was conducted up to 48 hours prior to the VR session in participants’ own time and location to encourage privacy and honest responding to sensitive questions (Gnambs & Kaspar, 2015), whilst keeping responses recent enough to reflect current conditions. Item order within all measures was randomised to minimise response errors (Bowling, 2005).

Upon arrival at a 25-minute session, participants were briefed on the risk of cybersickness and their freedom to withdraw at any point. They were oriented with the HMD and controllers and fitted the equipment with assistance. A brief familiarisation trial was conducted to ensure participants could perform the necessary actions; this was streamed to the researcher’s laptop for confirmation. Following the training trial, participants removed the HMD and were checked for any signs of cybersickness or discomfort.

Participants then re-entered the VE and were instructed to walk physically to the table and use the paintbrushes and pigments to paint on the canvas for 6 minutes, which has been demonstrated as sufficient time to induce presence while minimising boredom effects (Zhang et al., 2018). Participants were instructed not to step more than one pace from the table to prevent contact with physical room boundaries, which can disrupt spatial presence (Slater, 2009). Immediately following the VR exposure, participants completed the post-exposure questionnaire on the researcher’s laptop, including measures of spatial presence and attention allocation, to maximise accuracy in retrospective reporting (Tourangeau et al., 2000).

### 2.5 Data Analysis

Univariate outliers were detected using the interquartile range (IQR) method and corrected to the variable median to retain statistical power (Mowbray et al., 2018); two outliers were corrected in attention allocation and two in spatial presence. Multivariate outliers were assessed using Mahalanobis distance and Cook’s distance; no influential cases exceeded standard thresholds and all were retained. Following outlier correction, all variables were standardised to z-scores prior to model entry. Distributional analyses confirmed that skewness was within acceptable limits (skewness/SE ratios ≤ |2.11|) for the primary analysis; a sensitivity analysis applying Yeo–Johnson power transformations was also conducted to assess robustness (see Section 3.3).

The primary mediation hypothesis was tested using path analysis within a structural equation modelling (SEM) framework implemented in lavaan (Rosseel, 2012), with bootstrap standard errors (1,000 samples). This approach is equivalent to Hayes’s (2022) PROCESS Model 4 for mediation testing. The indirect effect was evaluated using bias-corrected accelerated (BCa) bootstrap confidence intervals (2,000 samples). Post-hoc analyses examined dose-response effects by splitting the sample into tertiles based on TAS scores and testing the attention–presence relationship within each group.

## 3. Results

### 3.1 Descriptive Statistics and Correlations

Descriptive statistics and zero-order correlations for the key study variables are presented in Table 1. All pairwise correlations among the three core variables were positive. The strongest zero-order correlation was between attention allocation and spatial presence (r = .70, p < .001). The correlations between trait absorption and both attention allocation (r = .18, p = .135) and spatial presence (r = .17, p = .163) were small and non-significant at the bivariate level, consistent with the hypothesis that absorption’s influence is indirect.

**Table 1.** Descriptive Statistics and Pearson Correlations for Key Variables (N = 70)

| Variable | *M* | *SD* | Range | 1. | 2. | 3. | 4. |
| --- | --- | --- | --- | --- | --- | --- | --- |
| 1. Trait Absorption | 21.04 | 7.65 | 4–34 | — |  |  |  |
| 2. Attention Allocation | 25.41 | 4.40 | 6–30 | .18 | — |  |  |
| 3. Spatial Presence | 47.74 | 8.55 | 12–60 | .17 | .70*** | — |  |
| 4. Depression | 16.97 | 10.29 | 1–44 | .30* | .01 | -.02 | — |

*Note.* $N = 70$ for all variables except Depression ( $N = 69$ ). Correlations are computed on raw (untransformed) scores.
\* $p < .05$ , \*\*\* $p < .001$ .

### 3.2 Primary Mediation Analysis

Path analysis provided strong support for the mediation hypothesis (H2). The results are summarised in Figure 1 and reported below using standardised coefficients from the primary (outlier-corrected, non-transformed) analysis.

**Figure 1.**
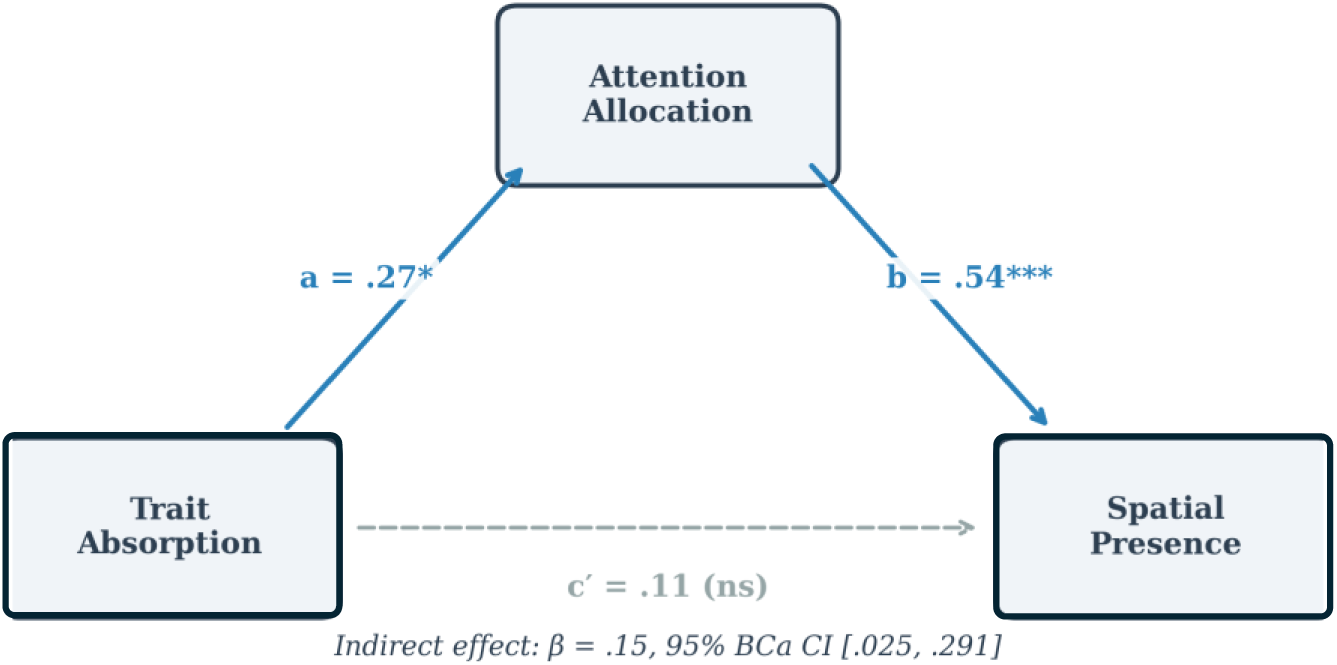
Path model of attention allocation mediating the relationship between trait absorption and spatial presence. Standardised path coefficients are shown. *** *p* < .001, * *p* < .05.

**Figure 2.**
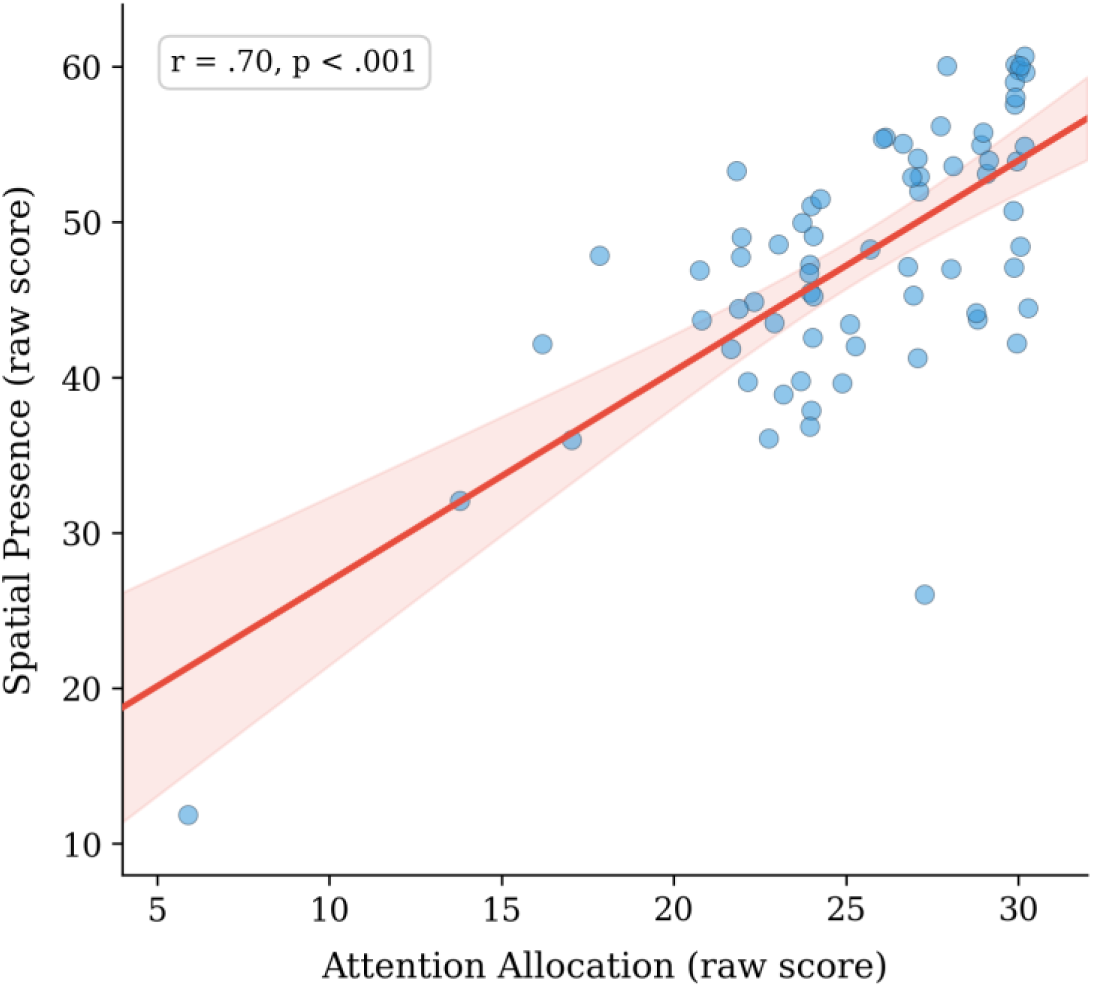
Scatterplot of attention allocation and spatial presence (raw scores, *N* = 70). The regression line with 95% confidence band is shown.

**Figure 3.**
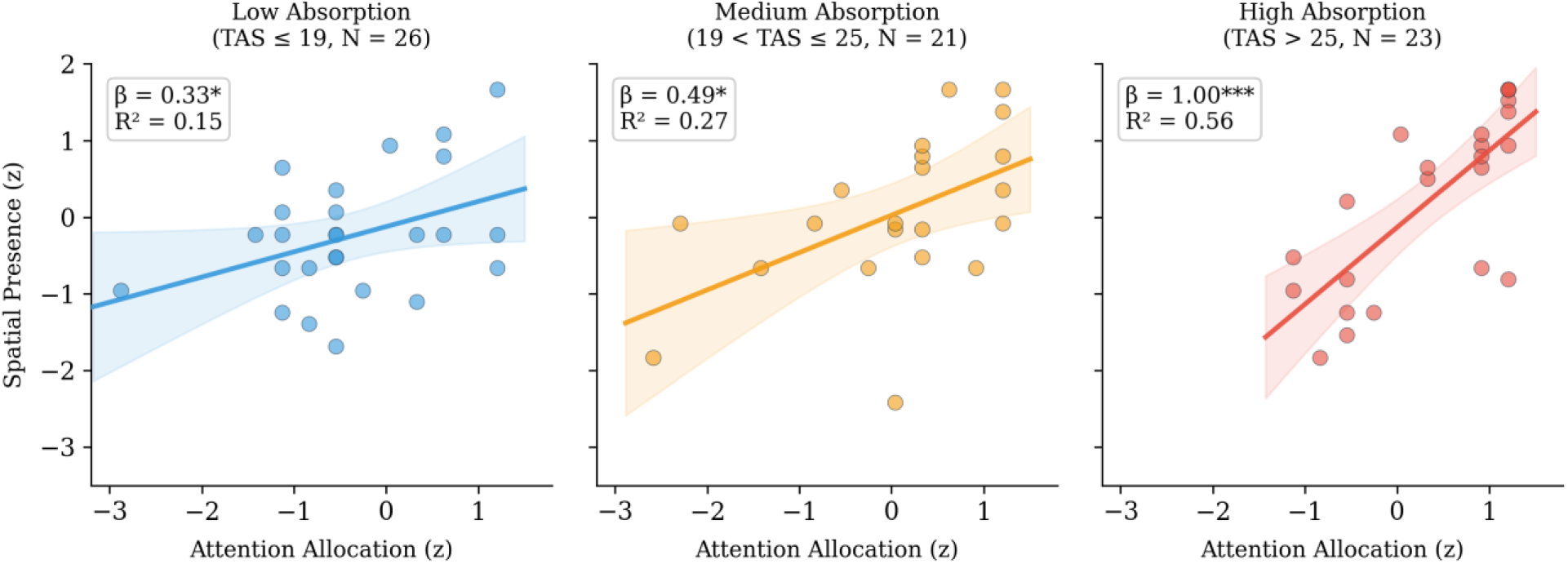
Dose-response relationship: scatterplots of attention allocation predicting spatial presence within each absorption tertile (outlier-corrected, standardised scores). Regression lines with 95% confidence bands are shown. The increasing slope across panels illustrates the amplification effect of trait absorption.

Path a (Absorption → Attention): Trait absorption was a significant positive predictor of attention allocation (β = 0.27, SE = 0.11, z = 2.50, p = .013, 95% CI [0.060, 0.494]). This confirms that higher trait absorption is associated with greater attentional engagement with the VE.

Path b (Attention → Presence): Attention allocation was a strong, significant positive predictor of spatial presence (β = 0.54, SE = 0.11, z = 4.95, p < .001, 95% CI [0.355, 0.786]). This was the strongest individual predictor in the model.

Direct effect, Path c’ (Absorption → Presence): After accounting for the mediation through attention, the direct path from trait absorption to spatial presence was no longer significant (β = 0.11, SE = 0.11, z = 0.98, p = .325, 95% CI [−0.124, 0.299]), indicating complete mediation.

Indirect effect: The indirect effect of trait absorption on spatial presence through attention allocation was statistically significant (β = 0.15, SE = 0.07, z = 2.22, p = .026, 95% BCa CI [0.025, 0.291]). This pattern of complete mediation shows that the influence of trait absorption on spatial presence is fully accounted for by its role in facilitating focused attention on the VE. The overall model accounted for 33.8% of the variance in spatial presence (R² = .338, Cohen’s f² = 0.51).

### 3.3 Robustness and Sensitivity Checks

Multiple analytical approaches confirmed the robustness of the mediation findings. First, robust regression (Yohai, 1987), which is less sensitive to outliers and distributional violations, yielded closely consistent estimates for the a-path (β = 0.28, p = .013) and for the effect of attention on presence in the full model (β = 0.55, p < .001; direct effect: β = 0.13, p = .281).

Second, because trait absorption and attention allocation exhibited mild residual negative skewness after outlier correction (skewness/SE ratios of −2.03 and −2.11, respectively), a sensitivity analysis was conducted in which Yeo–Johnson power transformations (Yeo & Johnson, 2000) were applied to all model variables prior to standardisation. This transformed analysis yielded substantively identical results: the indirect effect remained significant (β = 0.17; 95% BCa CI [0.033, 0.331]), and the model explained 35.7% of the variance in spatial presence (R² = .357, Cohen’s f² = 0.56). The convergence of results across untransformed and transformed analyses confirms that the mediation pathway is not dependent on distributional assumptions.

Third, tests for non-linearity (quadratic terms for absorption and attention) were all non-significant (all ps > .30), confirming that linear models appropriately characterise these relationships. Fourth, an exploratory moderation analysis found no evidence that depressive symptomatology moderated the attention–presence pathway (interaction β = −0.02, p = .874).

### 3.4 Post-Hoc Analyses: Dose-Response Relationship

To further characterise the role of trait absorption, a post-hoc dose-response analysis was conducted by splitting the sample into absorption tertiles based on TAS scores. The strength of the attention–presence relationship was then tested within each group. As shown in Table 2, the standardised regression coefficient for the prediction of spatial presence from attention allocation increased systematically across absorption groups. In the low-absorption group, attention explained 15.1% of the variance in spatial presence; this increased to 27.0% in the medium group, and reached 55.9% in the high-absorption group, where the regression coefficient approached unity (β = 1.00). This dose-response pattern indicates that trait absorption amplifies the effect of attentional focus on spatial presence.

**Table 2.** Dose-Response: Regression of Spatial Presence on Attention Allocation Across Absorption Tertiles.

| Absorption Group | *N* | Standardised $\beta$ | $R^2$ | *p* |
| --- | --- | --- | --- | --- |
| Low | 26 | .329* | .151 | .050 |
| Medium | 21 | .487** | .270 | .016 |
| High | 23 | 1.003*** | .559 | < .001 |

Note. $\beta$ = standardised regression coefficient for the prediction of spatial presence from attention allocation within each absorption tertile. \* $p < .05$ , \*\* $p < .01$ , \*\*\* $p < .001$ .

## 4. Discussion

This study investigated the psychological mechanisms linking individual differences to spatial presence in a highly immersive VE. Our findings support a complete mediation pathway, whereby trait absorption facilitates spatial presence entirely through its positive influence on attention allocation. Post-hoc analyses further uncovered a dose-response pattern, in which the strength of the attention–presence link systematically increased with higher levels of trait absorption.

### 4.1 Attention as the Complete Mediator

Contrary to claims that individual traits become negligible in highly immersive VEs (Sas & O’Hare, 2003; Uz-Bilgin & Thompson, 2022), the present well-powered study demonstrates that trait absorption remains a critical factor in shaping spatial presence. However, its influence is entirely indirect. Individuals higher in absorption are not simply ‘more present’; rather, their disposition enables them to direct attentional resources more effectively toward the VE, and it is this focused attention that directly builds the spatial presence experience. The finding that the direct path from absorption to presence was non-significant, while the indirect path through attention was robust, provides the first direct empirical support for the attentional mechanism proposed in foundational spatial presence models (Wirth et al., 2007; Hartmann et al., 2015).

This clarifies an important theoretical ambiguity. Although attention has been consistently emphasised as a prerequisite for spatial presence formation (Hofer et al., 2012; Schubert, 2009), and although absorption has been theorised to operate through attentional engagement (Wirth et al., 2007), the present study is, to the best of our knowledge, the first to empirically demonstrate this complete mediating pathway in a highly immersive VE. The complete mediation pattern indicates that without a corresponding increase in attention, trait absorption alone is insufficient to generate spatial presence.

These findings also help reconcile prior inconsistencies in the literature. The null effects reported for absorption in highly immersive contexts (Murray et al., 2007; Chiquet et al., 2023) may reflect not a genuine absence of influence, but rather the indirect nature of that influence, which would be missed when only direct relationships are tested. Previous null findings may also be attributable to methodological limitations, including the use of psychometrically questionable measures and underpowered samples (Parsons et al., 2015). By using the TAS, a well-validated measure of the absorption construct, and the MEC-SPQ, which cleanly separates spatial presence from its antecedents, the present study provides a more rigorous test.

### 4.2 Trait Absorption as a Cognitive Amplifier

Perhaps the more striking result is the dose-response relationship between absorption and the attention–presence link. The predictive power of attention on spatial presence was substantially stronger in participants with high trait absorption (explaining 55.9% of the variance) compared to those with low absorption (15.1%). In the high-absorption group, the standardised regression coefficient approached unity (β = 1.00), indicating nearly perfect sensitivity to attentional focus. This pattern suggests that absorption does more than simply direct attention; it appears to modulate the quality or efficacy of that attention in generating a coherent spatial model.

This amplification effect can be interpreted through the lens of predictive coding models of presence (Seth et al., 2012). These models propose that presence arises when sensory input from a VE successfully matches the brain’s internal predictions about the environment. Trait absorption, with its emphasis on deep perceptual engagement and openness to alternative representations of reality (Tellegen & Atkinson, 1974), may effectively lower the threshold for what constitutes a satisfactory ‘match’ between predicted and received sensory input. For high-absorption individuals, attentional focus may more readily translate into a coherent and convincing experience of the VE, effectively suppressing minor perceptual discrepancies between the virtual and the physical environment. For low-absorption individuals, the same level of attention may be less effective at bridging these mismatches, resulting in a weaker sense of presence.

This interpretation is consistent with earlier findings that trait absorption influences individuals’ judgements of VE realism and vividness of sensory experience (Baños et al., 1999), and with more recent conceptualisations of perceived realism as a separate but crucial contributor to presence formation (Weber et al., 2021). It also aligns with proposals that trait absorption’s motivational openness may suppress perceptual discrepancies between user expectations and system output (Chiquet et al., 2023), thereby ‘bridging the gap’ between imperfect technological input and a coherent presence experience, a gap widely demonstrated to reduce spatial presence (Fu et al., 2023; Slater et al., 1995).

### 4.3 Limitations

This study has limitations that qualify the conclusions drawn. First, the correlational, cross-sectional design precludes definitive causal claims; the mediation model represents the most theoretically plausible ordering of variables, supported by the data pattern, rather than confirmed causality. Second, the university student sample may limit generalisability, particularly to clinical populations. Third, although the overall model was well-powered for detecting the observed large effects (Cohen’s f² = 0.51), the sample of 70 limits the power to detect smaller effects, particularly in subgroup and moderation analyses. Fourth, the dose-response analysis was post-hoc and exploratory, requiring pre-registered replication. Fifth, all measures were self-report; future research incorporating physiological or behavioural indices of attention would strengthen causal inferences. Sixth, the largely non-clinical sample precluded a strong test of whether depression disrupts the attention–presence pathway; clinical samples with greater variance in depressive symptom severity would be needed to resolve this question. Finally, the VE used was a single, emotionally neutral environment, and generalisability to emotionally salient or therapeutically relevant contexts remains to be established.

### 4.4 Future Directions and Practical Implications

If trait absorption amplifies the translation of attention into spatial presence, it may be possible to design VR experiences that ‘prime’ an absorptive state, for instance through narrative framing or guided imagery that encourages imaginative engagement prior to the main experience. This could be particularly valuable for users who are naturally lower in trait absorption, potentially boosting the efficacy of VR-based therapy and training. Future research may also wish to test the separate dimensions of the TAS to clarify which facet(s) of absorption drive the amplification effect, as this would inform the design of targeted priming interventions.

Experimental designs that manipulate attention directly, such as dual-task paradigms or attentional cueing, would provide a stronger test of the causal role of attention. Similarly, comparing the mediating pathway across varying levels of system immersion would help clarify whether the mechanism identified here operates differently as a function of technological sophistication. Given the growing use of VR in clinical settings (Zlomuzica et al., 2024), extending this work to clinical samples remains a priority.

### 4.5 Conclusion

This study clarifies and extends our understanding of the individual factors that shape spatial presence in virtual reality. Attention allocation fully mediates the relationship between trait absorption and spatial presence in a highly immersive VE, and trait absorption acts as a cognitive amplifier, systematically strengthening the power of attention to create a convincing sense of ‘being there.’ Individual differences do not disappear in the face of powerful technology; they interact with it in consequential ways, shaping the quality of virtual experience. These findings carry implications for models of presence and for the design of VR-based interventions.

## Acknowledgements

The authors thank Jinyi Wang and Jumanah Elgamal who contributed to data collection, and the University of Leeds HELIX Digital Education Service for the loan of VR equipment. This research was conducted with ethical approval from the University of Leeds School of Psychology Research Ethics Committee (PSCETHS-1384).

## Author Contributions

HHH designed the study, collected data, conducted preliminary analyses, and contributed to manuscript preparation. CC supervised the research, conducted advanced statistical analyses, and prepared the manuscript. Both authors approved the final version.

